# The origin and structural evolution of *de novo* genes in *Drosophila*

**DOI:** 10.1101/2023.03.13.532420

**Authors:** Junhui Peng, Li Zhao

**Affiliations:** Laboratory of Evolutionary Genetics and Genomics, The Rockefeller University, New York, NY 10065, USA

**Keywords:** *De novo* genes, Drosophila, origin, structures, fold, ancestral sequence reconstruction, molecular dynamics

## Abstract

Although previously thought to be unlikely, recent studies have shown that *de novo* gene origination from previously non-genic sequences is a relatively common mechanism for gene innovation in many species and taxa. These young genes provide a unique set of candidates to study the structural and functional origination of proteins. However, our understanding of their protein structures and how these structures originate and evolve are still limited, due to a lack of systematic studies. Here, we combined high-quality base-level whole genome alignments, bioinformatic analysis, and computational structure modeling to study the origination, evolution, and protein structure of lineage-specific *de novo* genes. We identified 555 *de novo* gene candidates in *D. melanogaster* that originated within the *Drosophilinae* lineage. We found a gradual shift in sequence composition, evolutionary rates, and expression patterns with their gene ages, which indicates possible gradual shifts or adaptations of their functions. Surprisingly, we found little overall protein structural changes for *de novo* genes in the *Drosophilinae* lineage. Using Alphafold2, ESMFold, and molecular dynamics, we identified a number of *de novo* gene candidates with protein products that are potentially well-folded, many of which are more likely to contain transmembrane and signal proteins compared to other annotated protein-coding genes. Using ancestral sequence reconstruction, we found that most potentially well-folded proteins are often born folded. Interestingly, we observed one case where disordered ancestral proteins become ordered within a relatively short evolutionary time. Single-cell RNA-seq analysis in testis showed that although most *de novo* genes are enriched in spermatocytes, several young *de novo* genes are biased in the early spermatogenesis stage, indicating potentially important but less emphasized roles of early germline cells in the *de novo* gene origination in testis. This study provides a systematic overview of the origin, evolution, and structural changes of *Drosophilinae*-specific *de novo* genes.

## Introduction

*De novo* genes are novel genes born from scratch from previously non-genic DNA sequences (Begun et al., 2006; Levine et al., 2006; McLysaght & Hurst, 2016). Recent works support the existence of a considerable number of young *de novo* genes across various species and taxa, including humans and *Drosophila* (Begun et al., 2007; Cai et al., 2008; Chen et al., 2010; Heames et al., 2020; Heinen et al., 2009; Knowles & McLysaght, 2009; Li et al., 2010; Van Oss & Carvunis, 2019; L. Zhang et al., 2019; Zhao et al., 2014; Zheng & Zhao, 2022). While some studies proposed that proteins encoded by *de novo* genes tend to be highly disordered (Wilson et al., 2017) in order to prevent misfolding or aggregation, which can be neurotoxic in complex eukaryotes (Wilson et al., 2017), other studies suggested that *de novo* genes may not necessarily be disordered (Bornberg-Bauer et al., 2021; Bungard et al., 2017; Lange et al., 2021) and instead suggest that their structures could be highly conserved after their origination.

Despite the aforementioned advancements, our understanding of the protein structures of *de novo* genes remains very limited. It is still unclear whether *de novo* genes are capable of being well-folded, how frequently they are well-folded, and if they possess novel structural folds. The main obstacle was the lack of accurate, efficient, and scalable structural characterization tools that could be applied to a large amount of *de novo* genes. Here, we applied AlphaFold2 (Jumper et al., 2021) computational predictions as well as ESMFold (Lin et al., 2023) to evaluate the foldability of *de novo* genes. With the rapid development in genome sequencing, sequence alignment, and deep learning techniques, AlphaFold2, along with other neural network approaches, e.g., trRosetta (J. Yang et al., 2020), RoseTTAFold (Baek et al., 2021), and ESMFold (Lin et al., 2023) have demonstrated the ability to predict protein structures with near-atomic accuracy. AlphaFold2 has been applied at genomic scales to predict protein structures of the human proteome and the proteomes of several other species (Tunyasuvunakool et al., 2021; Varadi et al., 2021). Although AlphaFold2 has been proved to be highly accurate, it predicts only a single static protein structure per protein sequence (Lane, 2023), which could hinder our understanding of the protein structures of *de novo* genes since proteins can be highly dynamic in cells. Molecular dynamics (MD) simulation has shown to be a valuable tool to investigate protein dynamics (Dror et al., 2012), study protein structure stability (Childers & Daggett, 2017), and evaluate or refine predicted or designed protein structures (Heo & Feig, 2020; Schlick & Portillo-Ledesma, 2021). Thus, we further carried out large scale MD simulations to characterize the structural stability and dynamics of the predicted protein structures. In addition, the increasing power of bioinformatic and computational approaches has made it possible to obtain highly accurate general structural properties of genes, including intrinsic structural disorder (Necci et al., 2021), relative solvent accessibility (S. Wang et al., 2016), and the probability of being transmembrane proteins (S. Wang et al., 2016) or signal proteins (Almagro Armenteros et al., 2019).

In addition to the structure of *de novo* genes, the evolution of their sequences and structures after origination remains unclear. To address this question, it is necessary to identify and compare branch-specific *de novo* genes ranging from very young and old within a relatively diverged lineage. However, due to low genome sequencing quality and high genome recombination rates in some species, this process could be very difficult, especially with increased divergence time (Vakirlis, Carvunis, et al., 2020; Van Oss & Carvunis, 2019). This limitation greatly hampered our understanding of the origination of *de novo* genes and how their sequences and structures evolve after origination. In addition, it has been historically difficult to distinguish between rapidly evolving genes and *de novo* originated genes (Moyers & Zhang, 2016; Weisman et al., 2020). However, with the recent advancement in whole-genome sequence alignments and its ability to progressively align the genomes of an entire phylogenetic tree, including diverged species with high accuracy (Armstrong et al., 2020), we can now identify *de novo* protein-coding gene candidates with high confidence through the support of synteny-based alignments and non-coding sequences in outgroup species.

In this work, we utilized progressive whole-genome alignments and multiple homology detection methods to identify 555 *de novo* protein-coding gene candidates in *D. melanogaster* that were born within the last ∼67 million years since the *Drosophilinae* lineage (Kumar et al., 2017) with the support of ortholog non-coding DNA sequences in their corresponding outgroup species. We performed bioinformatic analysis and computational structural modeling to predict the structural and functional properties of each *de novo* gene candidate. Additionally, we employed ancestral sequence reconstruction (ASR) to examine the sequence and structural evolution of these genes throughout their evolutionary history. Our results provided a systematic overview of the foldability of proteins encoded by *de novo* genes and the patterns in which their sequences and structures evolve after origination.

## Results

### Identification of *Drosophilinae* lineage-specific *de novo* gene candidates in *D. melanogaster*

We built the whole genome alignment of 20 *Acalyptratae* fly species (Figure 1A) using Progressive Cactus Aligner (Armstrong et al., 2020). A summary of the alignments can be found in Figure S1. Overall, the protein-coding bases in *D. melanogaster* can be aligned to closely related species at high coverage (97% with *D. sechellia*), while the coverage dropped significantly to around 50% in relatively distant species (49% with *B. dorsalis*). For the 13968 annotated protein-coding genes in *D. melanogaster* investigated in our study, we observed 13798 (98.8%) of them were covered in cactus alignments. For each of the 13798 *D. melanogaster* protein-coding genes, we combined homology obtained from all-vs-all *blastp* (Altschul et al., 1990) analysis, *Genewise* (Birney et al., 2004) and *Spaln* (Iwata & Gotoh, 2012) predictions to identify annotated/unannotated orthologs and non-genic hits from their syntenic regions (Figure 1B). For simplicity, we termed the furthest branches that have annotated/unannotated orthologs as *Br_i_*, where *i* could range from 1 to 9 for each potential candidate, as shown in Figure 1A. The above step gave 1285 potential *de novo* gene candidates within *Br_9_* (see Figure 1B and *Material and Methods* for detail, *supplementary File S1: unannotated syntenic regions*). We then removed genes that have homologs that are not in the syntenic regions using all-vs-all *blastp* (Altschul et al., 1990). This led to 686 potential *de novo* gene candidates within *Br_9_*. As the last filtering step, we removed candidates that have reliable annotated or unannotated homologs outside of *Br_i_* by *blastp* and iterative *jackhmmer* (S. R. Eddy, 2011) search against UniProt Knowledgebase sequence database (UniprotKB) (Bateman et al., 2022) and *tblastn* search against NCBI representative genomes (Figure 1B, *Material and Methods*). Finally, combined with homology and synteny, we identified 555 *de novo* protein-coding gene candidates in *D. melanogaster* that potentially originated within *Drosophilinae* lineage (*supplementary File S2: The list of de novo gene candidates and their properties*). All these *de novo* gene candidates were supported by evidence of possible ancestral noncoding DNA sequences. For each of the branches (*Br_1_*, *Br_2_* to *Br_9_*), the number of *de novo* gene candidates originated in the branch is shown in Figure S2. We did not observe *de novo* genes that are *D. melanogaster* specific due to the identification of putative unannotated “partial” orthologs (*Material and Methods*) in unannotated syntenic regions in the outgroups. However, we did observe 73 *D. melanogaster* protein-coding genes with unannotated syntenic regions in *D. simulans*, *D. sechellia,* or other species in more distant branches. Of these 73 genes, 50 were not considered to be *de novo* genes since the unannotated syntenic regions were predicted to be unannotated orthologs (*Material and Methods*) with high confidence. The remaining 23 genes were identified as *de novo* gene candidates, but were inferred to have originated from more distant branches. Based on the location of ancestral noncoding DNA sequences, we found that 397 of them were born from intergenic regions, and 158 from intragenic regions.

**Figure 1.**
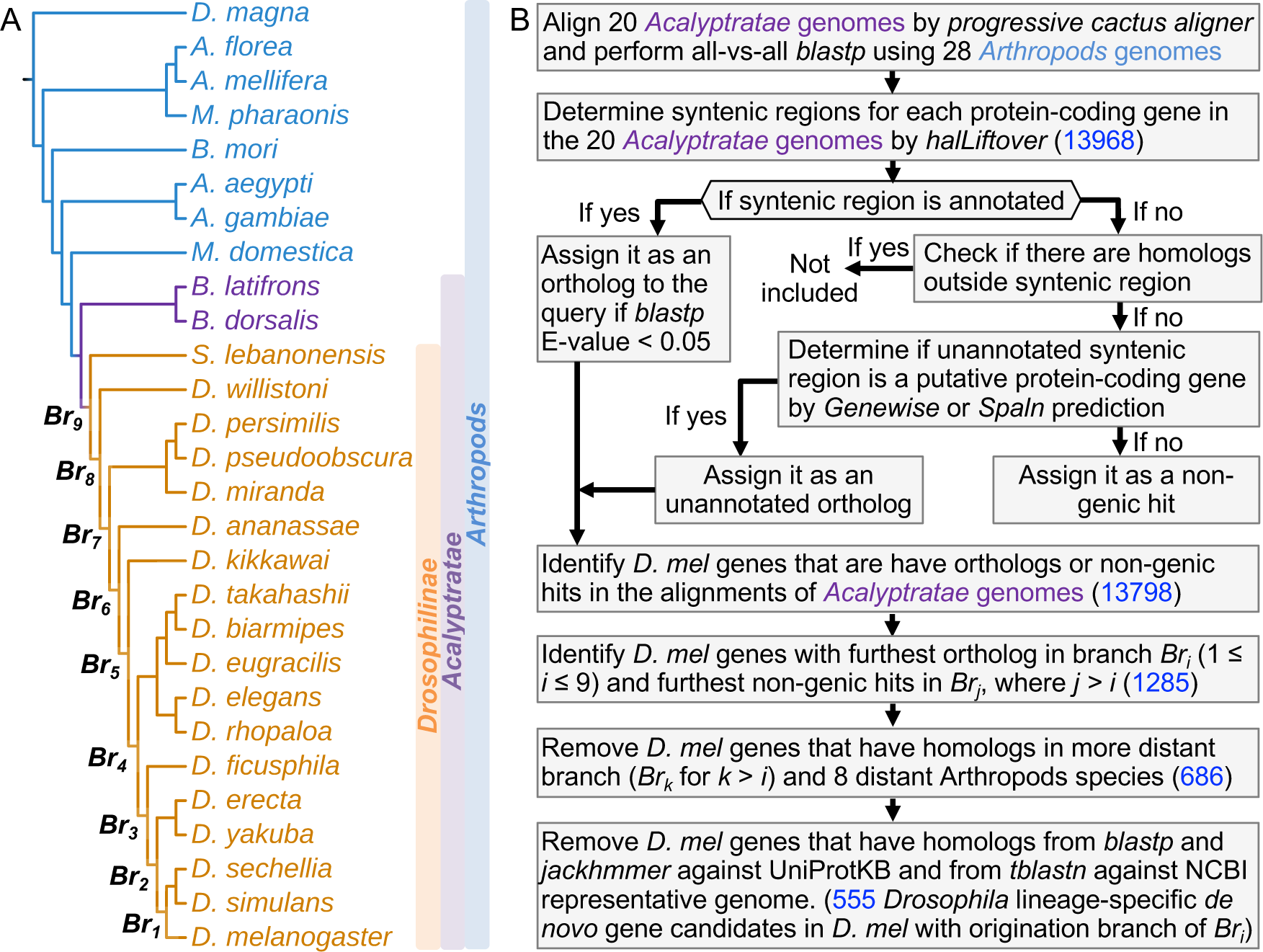
Workflow to identify *Drosophillinae* lineage-specific *de novo* genes in *D. melanogaster*. (A) Phylogenetic tree of the *Drosophilinae*, *Acalyptratae*, and *Arthropods* species. Progressive cactus genome alignment was performed among the 20 *Acalyptratae* genomes. (B) The workflow and pipeline to identify *Drosophilinae* lineage-specific *de novo* genes and track the origination within the *Drosophilinae* lineage for the current protein-coding genes in *D. melanogaster* (see *Material and Methods*). The numbers of potential candidates in each identification step were highlighted in blue.

### The origin of *Drosophilinae*-specific *de novo* genes is more likely to be associated with open chromatins than transposable elements

Our analysis shows that most *de novo* gene candidates had biased expression in the testis, head, and ovary. Thus, to understand their relationships with open chromatin regions, which are enriched with regulatory sequences, we examined ATAC-seq data from the three tissues (Witt et al., 2021). We found that both intergenic and intragenic *de novo* gene candidates have more peaks in their nearby ±500 bp regions than putative random ORFs in intergenic regions. In these regions, the peaks of the *de novo* gene candidates also have higher peak intensities. When considering broader regions, we found that *de novo* gene candidates, regardless of being intergenic or intragenic, were closer to their nearest peaks. This suggests that *de novo* genes are partly associated with open chromatin conformation changes. To assess the potential involvement of TEs in *de novo* gene origination, we searched for DNA repeat signals (see Material and Methods) in these regions. However, we did not find evidence of an association between *de novo* gene candidates and TEs in *Drosophila*, contrasting with previous reports of up to 20% of *de novo* transcripts being associated with TEs in primates, including humans (Ruiz-Orera et al., 2015). One possible explanation is that the primate genomes have much higher TE contents than fruit flies; thus the potential regulatory roles of TEs in *Drosophila* new genes are limited. Our findings suggest that in *Drosophila*, lineage-specific *de novo* genes are associated with open chromatin regions (Figure S3) but not TEs, a different pattern from what has been observed in primates.

### *De novo* gene candidates are mostly adaptive and shaped by both adaptive and non-adaptive changes

Compared to other annotated genes, *de novo* gene candidates display distinct properties in sequence composition (GC content), sequence evolution (ω, ω_a_, ω_na_, and α), structural properties, and expression patterns (Figure 2). Specifically, these candidates exhibit lower GC contents (Figure 2A), and this trend applies to the codons of each amino acid that contains G or C in their codons (Figure 2B). We further computed the optimized codons for each amino acid within the *D. melanogaster* genome (*Material and Methods*). We found that *de novo* genes tend to use less optimized codons than other protein-coding genes (Table S1). These findings suggest that *de novo* genes might be using unoptimized codons and that selection on codon usage might be playing important roles in their evolution.

**Figure 2.**
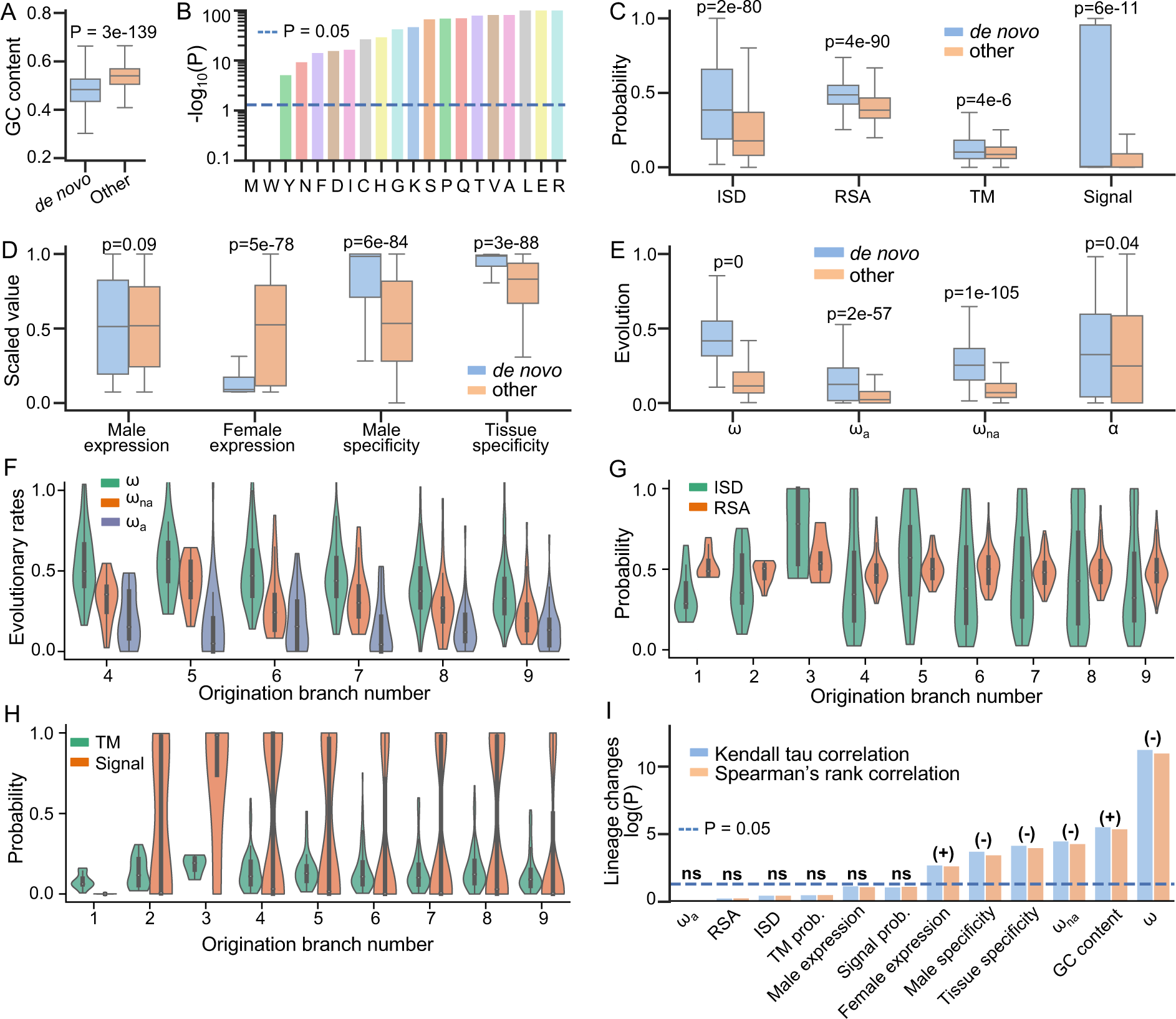
Special properties of *de novo* gene candidates. (A) GC contents of *de novo* gene candidates (denoted as “*de novo*”) were significantly lower than other annotated protein-coding genes (denoted as “other”). (B) GC contents of the codons utilized by each of the amino acids in *de novo* gene candidates were significantly lower than those of the amino acids in other annotated protein-coding genes. *De novo* gene candidates also differed in structural properties (C), expression patterns (D), and evolutionary patterns (E). (F) Evolutionary rates of *de novo* gene candidates with different gene ages. Due to insufficient data, *Br_1_*, *Br_2_*, and *Br_3_* were not shown. (G) Structural disorder (ISD) and solvent accessibility of de novo gene candidates with different gene ages. (H) The probability of containing transmembrane or signal proteins for *de novo* gene candidates with different gene ages. (I) Correlations between different properties of *de novo* gene candidates and their gene ages. Positive correlations and negative correlations were indicated by (+) and (−) respectively. Expression patterns and evolutionary patterns of *de novo* gene candidates were significantly correlated with gene ages, while structural properties were not.

Compared to other protein-coding genes, the protein products of *de novo* genes tend to be more disordered (p=2e-80), more exposed (p=4e-90), and are more likely to be transmembrane proteins (p=4e-6) or secretory proteins (p=6e-11) (Figure 2C). *De novo* genes have higher male specificity (p=6e-84), higher tissue specificity (p=3e-88), relatively lower expression levels in females (p=5e-78), and slightly higher but not significant expression levels in males (p=0.08). Many of these patterns are consistent with observations in studies that were focused on less divergent species groups in *Drosophila* or other taxonomy (Begun et al., 2007; Heames et al., 2020; Palmieri et al., 2014; Vakirlis et al., 2018; Zhao et al., 2014), further indicating that *de novo* genes exhibit some universal patterns that are not dependent on their lineage or identification method.

By analyzing comparative and population genomics data, we found that *de novo* genes are under faster sequence evolution compared to other protein-coding genes (Figure 2E). The adaptation rates, nonadaptation rates, and proportions of adaptive changes of *de novo* genes are higher than other genes (Figure 2E), indicating that both adaptive evolution and relaxation of purifying selection contribute to the elevated evolutionary rates of *de novo* genes. The results that *de novo* genes are more likely to be transmembrane proteins or secretory proteins (Figure 2C) and male-specific or tissue-specific (Figure 2E) suggest that some *de novo* genes might have specific molecular or cellular function (Lange et al., 2021; Rivard et al., 2021; Vakirlis, Acar, et al., 2020).

### *De novo* gene candidates undergo gradual sequence/function changes without significant structural changes

To understand whether and how structures change after the fixation of a *de novo* protein-coding gene, we then studied structural differences in *de novo* genes among different origination branches or gene ages. We compared the differences in protein properties of *de novo* genes with origination branches (*Br_1_* to *Br_9_*, Figure 1A) by applying Kendall tau and Spearman’s rank correlation analysis. We observed significant sequence changes among different origination branches. For example, the sequence evolutionary rates (ω) of *de novo* genes are significantly negatively correlated with origination branches (Kendall tau P = 1e-11, Spearman’s rank P = 5e-12, Figure 2F and 2I). The significant changes of ω might be dominated by the decrease of non-adaptive changes (ω_na_) since non-adaptation rates had significant negative correlations with origination branches as well (Kendall tau P = 5e-5, Spearman’s rank P = 3e-5, Figure 2F and 2I), while the negative correlations between adaptation rates and origination branches were not significant (Kendall tau P = 0.9, Spearman’s rank P = 0.9, Figure 2F and 2I). The decrease of non-adaptive changes (ω_na_) might indicate an increase in the strength of purifying selection, suggesting the functional importance of *de novo* genes increase over limited evolutionary time.

We found that GC contents are lower for *de novo* genes originated in younger branches and lower in older branches (Kendall tau P = 4e-6, Spearman’s rank P = 3e-6, Figure 2I). The same trend applies to the GC content of the codons of each amino acid (Table S1). We computed the optimal codon, most frequently used codon, for each amino acid. We found that younger *de novo* genes used significantly less optimal codons than older *de novo* genes (Table S1). These observations might suggest an important role of selection on codon usage or translation in the evolution of *de novo* genes. We observed significant correlations between male specificity and tissue specificity and origination branches (Kendall tau P = 3e-4 and 2e-4, Spearman’s rank P = 1e-4 and 7e-5, Figure 2I). Similarly, we observed weaker but significant correlations for female expression levels (Kendall tau P = 2e-3 and 2e-3), and weaker but not significant correlations for male expression levels (Spearman’s rank P = 0.07 and 0.06, Figure 2I). The correlations between expression patterns and origination branches might indicate a gradual change in protein functions, which further suggest that these *de novo* genes are under certain degrees of selection.

Interestingly, we did not observe significant changes in structural properties among different origination branches, including structural disorder (ISD), solvent accessibility, probability of being transmembrane proteins, and probability of being signal proteins from Kendall tau and Spearman’s rank test (Figure 2G, 2H, and 2I). This indicates that upon origination, the overall structural properties of *de novo* genes might remain similar in *Drosophilinae* lineage. The correlations between *de novo* gene properties and their gene ages indicate that these *de novo* gene candidates might undergo gradual sequence and functional changes without significant structural changes.

### A small subset of *de novo* genes is potentially well-folded with complex structural folds

We used AlphaFold2 (Jumper et al., 2021) to predict the structural models of *de novo* gene candidates. We further used the per-residue confidence score (pLDDT) from AlphaFold2 predictions to estimate the foldability of these candidates as high pLDDT scores often indicate accurate protein folding, and low pLDDT scores highly correlate with protein disorder (Jumper et al., 2021). We showed that pLDDT from AlphaFold2 predictions strongly correlated with pLDDT from RoseTTAFold (Baek et al., 2021) predictions (Pearson correlation R = 0.77, P = 1e-43), ESMFold (Lin et al., 2023) predictions (Pearson correlation R = 0.76, P = 2e-42), and the convergence of trRosetta (J. Yang et al., 2020) predictions (Pearson correlation R = 0.60, P = 2e-22) (Figure 3A, Material and Methods), suggesting consistency among the three state-of-the-art prediction methods. We categorized *de novo* gene candidates into three structural groups, each for 1) potentially well-folded, 2) partially folded, and 3) not folded. We defined genes to be potentially well folded if their average per-residue confidence score (pLDDT) were greater than 0.8 and the percent of confidently predicted residues (pLDDT > 0.7) greater than 90%; genes to be potentially partially folded if more than 30% of their residues or more than 50 consecutive residues being confidently predicted (pLDDT > 0.7); and the remaining genes to be potentially not folded. We found that most of the *de novo* gene candidates might only be partially folded (297/555) or not folded (224/555) (Figure 3B, *supplementary File S2: The list of de novo gene candidates and their properties*).

**Figure 3.**
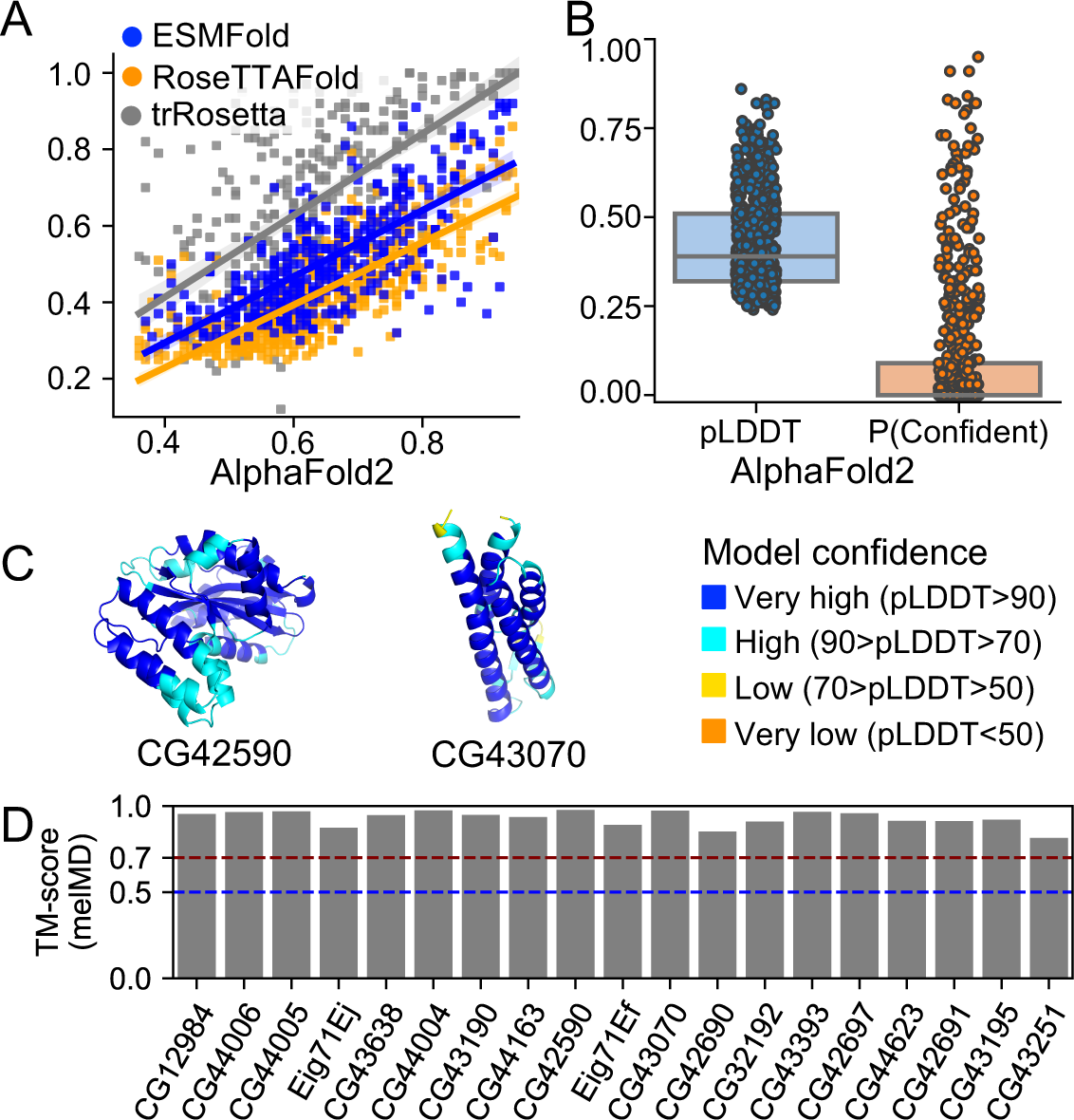
AlphaFold2 structure predictions of *de novo* gene candidates. (A) AlphaFold2 predictions are strongly correlated with RoseTTAFold and trRosetta predictions. (B) Most *de novo* gene candidates might not have the ability to be well-folded according to pLDDT, per-residue confidence score, and P(confident), the percent of residues that were confidently predicted. (C) Two examples of potentially well-folded *de novo* gene candidates: CG42590 and CG43070. In both cases, the core regions were predicted with very high confidence (blue). (D) The predicted structural models of the 19 potentially well-folded *de novo* genes retained highly similar structural folds during MD simulations with pairwise TM-score of representative structures larger than 0.70.

Interestingly, we found 35 *de novo* gene candidates that are potentially well-folded. Among these 35 *de novo* genes, 16 might only fold into simple folds containing a single or two α-helices, while another 19 might fold into complex folds (Table S2). We further performed 200 ns MD simulations for each of the 19 structural models to refine the structural models and characterize the stabilities of the structural folds (see Material and Methods, Table S3). The structural models all retained highly similar structural folds during the MD simulations, with pairwise TM-scores of representative conformations larger than 0.70 (Figure 3D). For example, CG42590 and CG43070 had averaged pLDDT scores of 0.90 and 0.89 (Figure 3C, Table S2). Their structural folds remained highly stable during MD simulations with averaged pairwise TM-score values of 0.98 and 0.92 (Table S3).

### Most potentially well folded *de novo* genes adopt existing protein structure folds

To check if potentially well-folded *de novo* genes have novel structural folds, we compared their MD-refined structures with all experimentally determined protein structures in protein data bank (PDB) (Berman, 2000). We did this by searching against PDB for potential novel structural folds using RUPEE (Ayoub & Lee, 2019). Interestingly, we found that most of them (16/19) have similar structural folds in PDB (Table S2). These similar structural folds were not due to sequence similarity. In contrast, the sequence identity between the *de novo* gene candidates and their similar structures was mostly less than 10% (Table S2). For example, the structural model of CG43195 is similar to the A chain of PDB structure 1U89 with a TM-score of 0.70. However, their sequence identity is only 4% (Table S2). Notably, we also found 3 *de novo* gene candidates, Eig71Ei, Eig71Ej, and CG43251, with maximum TM-score against protein structures in PDB smaller than 0.5 (Table S2, Figure S4), suggesting that the 3 candidates might adopt novel structural folds that have not been identified before. Overall, our results indicated that well-folded *de novo* genes are likely to adopt existing protein structure folds.

### Most potentially well-folded *de novo* genes are likely to be born well-folded

We showed that *de novo* gene candidates could undergo fast sequence adaptation without significant structural changes. We further investigate the structural changes of *de novo* genes after origination using the 19 potentially well-folded *de novo* genes as examples. For all the 19 *de novo* genes, we reconstructed their ancestral states, including most ancestral states and intermediate states in other branches, by ancestral sequence reconstruction (ASR). We then used AlphaFold2 to predict the 3D structures of these ancestral states. We found that the ancestral states of the 19 *de novo* genes were all predicted to be potentially folded at high confidence (average pLDDT > 0.7, Table S2). We further compared the structural models of ancestral states to current states. We found that they share similar structural folds with all pairwise TM-scores greater than 0.56 (Figure 4A, top panel), suggesting that these potentially well-folded *de novo* genes are likely to be born with similar structural folds to the current forms. To refine the structural models and characterize the stabilities of the structural folds, we further conducted 200 ns MD simulations starting from the ancestral structural models (see *Materials and Methods*, summarized in *supplementary File S3: MD simulations of ancestral state structural models*). We found that most of the ancestral state structural models retained similar structural folds to their current *D. melanogaster* forms during MD simulations, with almost all pairwise TM-scores of representative conformations close to or larger than 0.70 (Figure 4A, middle panel and bottom panel). In addition, we found that the structural fold stabilities and their similarities to their current forms did not change with the age of ancestral states (Figure 4B). Taking CG42590 as an example, MD simulations of CG42590 in its current form revealed that it contains a highly stable structural fold (Figure 4C, left column). Meanwhile, MD refined structural models of CG42590 ancestral states revealed that they are highly similar to CG42590 current form (Figure 4D, right column), with an average TM-score of 0.90. Interestingly, we also observed one exception, CG43251. We found that CG43251 might undergo substantial global structural changes from ancestral states to their current state (Figure 4A, last column), with an average TM-score of 0.32 to their current form. In the MD refined structure of the current form, CG43251 forms a stable fold with an alpha-helix in the N-terminal and a β-hairpin in the C-terminal (Figure 4D, left column), while in its ancestral states, the structures were disordered (Figure 4D, right column). Altogether, the results suggested that after origination, most potentially well-folded *de novo* genes might preserve a similar fold as they were born, while in some rare cases, the candidates undergo substantial structural changes.

**Figure 4.**
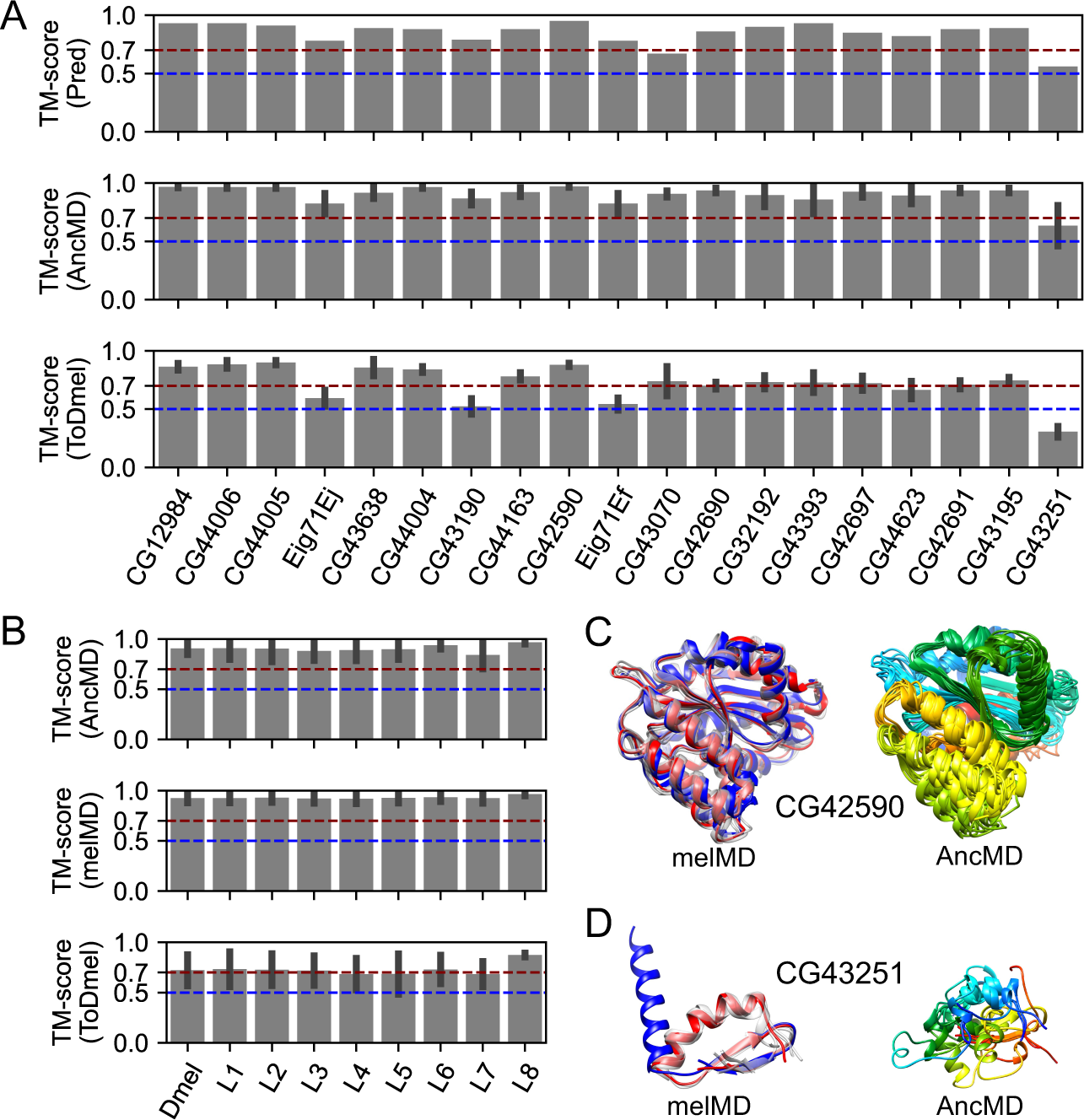
Structural evolution of 19 potentially well-folded *de novo* gene candidates. (A) Ancestral states have similar structural folds to their current forms as predicted by AlphaFold2 (“Pred”, top panel). The structural folds of ancestral structural models were stable (“AncMD”, middle panel) and similar to their current forms (“ToDmel”, bottom panel) during MD simulations, except for the case of CG43251 (last column). (B) Structural fold stability of ancestral states (“AncMD”, top panel), and current forms (“melMD”, middle panel) as a function of the ancestral state ages. Structural models of ancestral states were similar to their current forms, regardless of their ancestral branches (“ToDmel”, bottom panel). (C and D) Two examples of the structural evolution of potentially well-folded *de novo* gene candidates. AlphaFold2 predicted structural model (blue), MD refined structural model (red) and other representative structural models (gray) of CG42590 (C, left panel) and CG43251 (D, left panel). MD refined structural models of different ancestral states of CG42590 (C, right panel) and CG43251 (D, right panel) colored by residue index, with N-terminus being blue and C-terminus being red. Ancestral states of CG42590 have similar structural folds while those of GC43251 are different.

### Early germline cells in testis are non-negligible in *de novo* gene origination, despite that most *de novo* genes are enriched in later germ cells

Of the 555 *de novo* gene candidates identified, many of them (217, ∼40%) had biased expression in the testis. To investigate how *de novo* genes originated and evolved in testis, we analyzed the expression patterns of testis-biased *de novo* genes using testis single-cell RNA-sequencing data (Witt et al., 2019). We used the expression patterns of all annotated testis-biased *Drosophila melanogaster* genes to cluster these genes into four different clusters (Figure 5A, Figure S5). We numbered the four clusters according to the expression patterns, where genes in cluster #1 tend to be highly expressed in early spermatogenesis stage, genes in cluster #2 showed average expression in spermatogonia and spermatocytes stages, genes in cluster #3 showed average expression in spermatocytes and spermatids stages, and genes in cluster #4 showed peak expression in spermatids stage. We found that testis-biased *de novo* genes’ expression profile is similar to other testis-biased genes (Figure 5B). We then asked whether *de novo* genes in different clusters showed different structure/sequence characteristics. We found that *de novo* gene candidates in cluster #1 were significantly different from other genes. For example, these *de novo* gene candidates were less likely to be transmembrane or secretory proteins, and they were more likely to be disordered or exposed proteins than *de novo* gene candidates in other clusters (Figure 5C). We also found that, compared to other clusters, *de novo* genes in cluster #1 evolved faster, and the faster evolutionary rates were mostly contributed by their faster adaptation rates (Figure 5D). These special properties of *de novo* genes in cluster #1 were not because testis-biased genes in cluster #1 have these properties. We compared *de novo* genes candidates and other testis-biased genes in cluster #1. We found that in cluster #1, *de novo* genes were more likely to be disordered and exposed than other testis-biased genes, and that *de novo* genes tend to evolve faster with faster adaptation rates. In cluster #1, the faster evolutionary rates of *de novo* genes were also more likely to be contributed by their faster adaptation rates (Figure S6). While previous studies emphasized the importance of mid-to-late spermatogenesis stages in *de novo* gene origination (Kaessmann, 2010; Soumillon et al., 2013; Van Oss & Carvunis, 2019; Witt et al., 2019), our results indicate that although only very few *de novo* genes were enriched in the early spermatogenesis stages, they might also play a non-negligible role in *de novo* gene origination.

**Figure 5.**
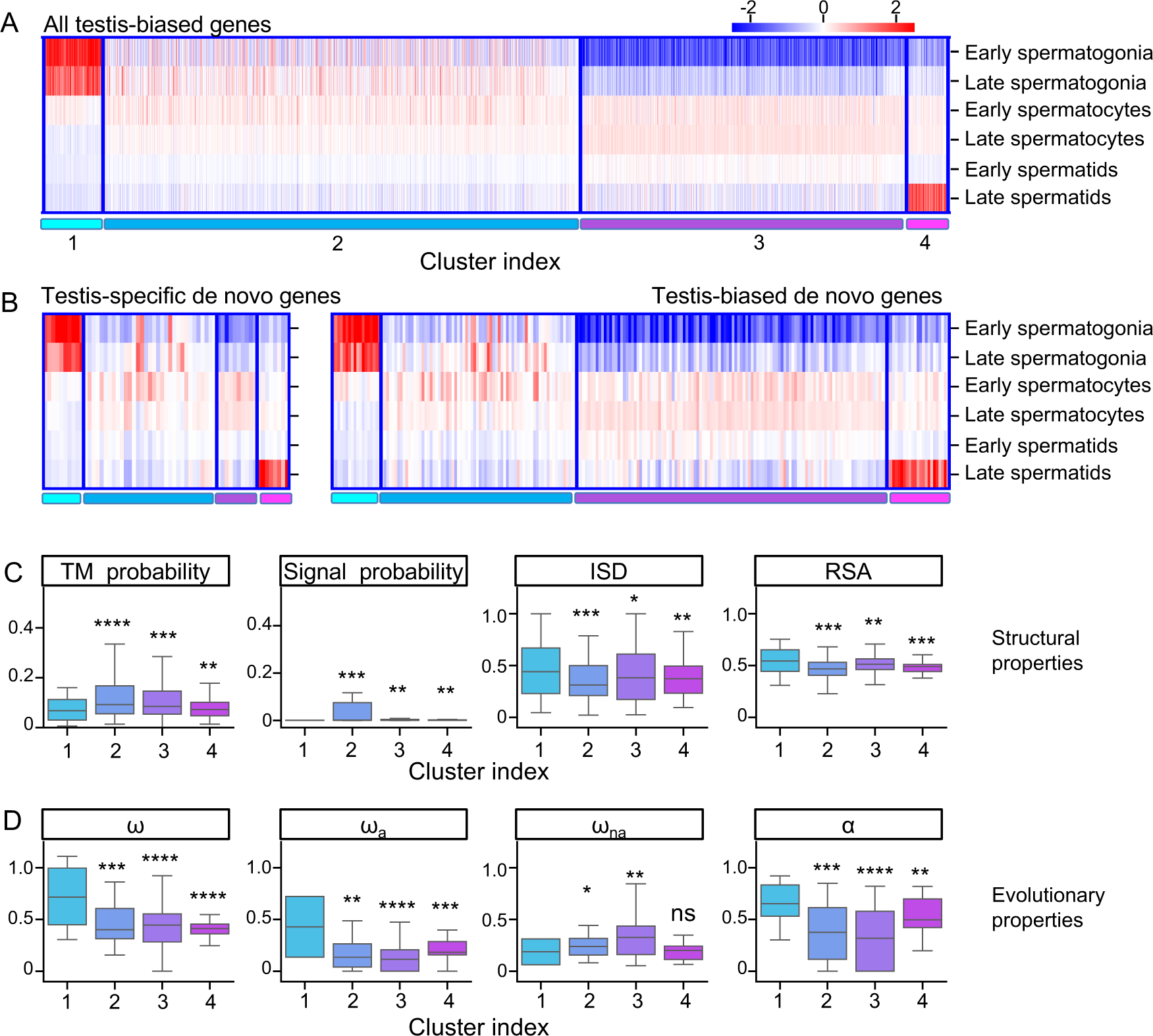
Expression pattern of testis-biased *de novo* gene candidates. (A) Clustering of all *D. melanogaster* testis-biased genes. (B) Expression patterns of testis-specific (left) and testis-biased (right) *de novo* gene candidates in different clusters. (C) *de novo* gene candidates in Cluster #1 are less likely to be transmembrane (TM probability) or signal (Signal probability) proteins but are more likely to have higher structural disorder (ISD) and higher solvent accessibility (RSA). (D) *de novo* gene candidates in Cluster #1 have higher evolutionary rates (ω), adaptation rates (ω_a_), and proportions of adaptive changes (α), but not higher nonadaptation rates (ω_na_).

## Discussion

In the past decade, various synteny-based methods have been applied to determine the *de novo* emergence of *de novo* genes (Van Oss & Carvunis, 2019). However, many of them suffered from divergence between genomes in longer timescales, fragmented genome assemblies, and high genome recombination rates (Van Oss & Carvunis, 2019). Thus, in this study, we used whole-genome alignments instead of synteny mapping to determine the ortholog regions of *D. melanogaster* protein-coding genes.

Specifically, we used the base-level progressive cactus aligner, which has been proved to be highly accurate in both pairwise genome alignments (Paten et al., 2011) and progressive genome alignments of an entire phylogenetic tree (Armstrong et al., 2020). With the high-quality base-level whole genome alignments, we can identify *de novo* genes with the support of alignments against non-coding sequences in outgroup species. The aligned non-coding sequences could also facilitate the discovery of possible origination events of *de novo* genes. Despite these merits, there are still some inherent drawbacks to this method. The first is that we did not include taxonomically restricted genes (TRG) or orphan genes that cannot be aligned in progressive cactus alignments, which could underestimate the number of true *de novo* originated genes. However, it is also important to note that, although TRG or orphan genes are likely to be *de novo* emerged, without the support of noncoding alignments, we cannot rule out the possibility of other mechanisms, such as horizontal gene transfer (HGT), transposition, etc. The second drawback is that we used simulated data to distinguish unannotated orthologs and unannotated noncoding regions. For genes with unannotated syntenic regions, we define orthology if their gene structure prediction scores were significantly beyond random expectations. In this case, unannotated ortholog regions with frameshift indels but high prediction scores might also be considered orthologs. Thus, it is likely to assign a number of *de novo* genes to a more ancestral branch than their true origin dates. However, we argue that, according to the high average genomic mutation rates in *Drosophila* species (Haag-Liautard et al., 2007), an unannotated ortholog region with high prediction scores might be under strong purifying selection or might just be dead recently, otherwise this region would be completely random after random neutral mutations for millions of years. This compromise might be necessary as misidentification of old genes as young genes may lead to biased patterns (Moyers & Zhang, 2015, 2016; Weisman et al., 2020). We further found gene expression evidence from five different *Drosophila* species for many of these unannotated regions (Figure S7). Note that it is still likely that we might have excluded potential genic sequences that were too diverged to have meaningful gene structure predictions (Iwata & Gotoh, 2012). Another drawback was that the estimation of the origin branch might be affected by the unavailability of high-quality genomic data for some other species in the branch of interest. With the great efforts being made to improve the genome sequencing and annotation qualities, we will be able to better study *de novo* gene emergence events in the future.

There were a few case studies trying to characterize the foldability and structure of proteins encoded by some specific *de novo* originated genes (Bornberg-Bauer et al., 2021; Bungard et al., 2017; Lange et al., 2021). However, it remains unclear whether *de novo* genes could be well-folded with complex structural folds, how often *de novo* genes could be well-folded and whether *de novo* genes could have novel structural folds. Here, we explored the foldability of *de novo* genes/proteins by highly accurate and efficient AlphaFold2 predictions. In general, the AlphaFold2 predictions correlated significantly with other state-of-the-art neural network approaches, such as trRosetta, RoseTTAFold, and ESMFold, and were comparable to high-resolution structure determination techniques, such as solution NMR and x-ray crystallography, especially for small proteins (Tejero et al., 2022). We found that most *de novo* genes might be partially folded or not folded, and only very few of them have the potential to be well-folded with complex structural folds (19/555), among which several (3/19) might have novel structural folds. The complex structural folds of these *de novo* gene candidates remained stable during 200 ns MD simulations (Figure 3D), which further suggested the high accuracy of AlphaFold2 predictions. Our results provided a systematic overview of the foldability of *Drosophilinae* lineage-specific *de novo* genes and highlighted the possibility that *de novo* proteins can be well-folded and even folded into novel protein folds. However, these observations were based on computational predictions. Further experimental validation would be needed to better understand the protein structures of *de novo* genes.

The fact that *de novo* genes were preferentially expressed in testis has led to the hypothesis that *de novo* genes were “out-of-testis”. The “out-of-testis” hypothesis was thought to be related to the facilitating role of the permissive chromatin state in germline cells (Majic & Payne, 2020). Similar findings were observed in our study - we found that *de novo* genes were associated with open chromatins in testis (Figure S3), and that more than half of the identified *de novo* gene candidates were biased toward testis. These findings highlight the important role of testis and chromatin state in the origination of *de novo* gene origination. In addition, an examination of another two hotspot-tissues, head, and ovary, suggested that *de novo* gene candidates were also associated with open chromatins in these two tissues (Figure S3), highlighting the non-neglectable role of open chromatins in the emergence of *de novo* genes. In agreement with previous findings that spermatocytes and spermatids were the “hotspots” for *de novo* genes (Witt et al., 2019), we found that most testis-biased *de novo* gene candidates identified here have their peak expression in late spermatogenesis stages, including spermatocytes and spermatids. The results might indicate that *de novo* genes that originated in late spermatogenesis stages may have higher probabilities of being fixed due to a stronger fitness effect in sperm competition, while those that originated in early spermatogenesis stages were more likely to be lost during evolution. On the other hand, we found that testis-biased *de novo* genes in early spermatogenesis stages showed some special properties compared to testis-biased *de novo* genes in later stages. For example, these genes tend to be more disordered or exposed and evolve faster with higher adaptation rates. Our results indicate a non-negligible role of early spermatogenesis stages in *de novo* gene origination. To further address the precise mechanisms, it would be necessary to conduct functional assays to examine the fitness effect of young and old *de novo* genes, which is beyond the current scope of this study.

It has been debated whether *de novo* genes change their protein structures after origination, and it is still unclear how *de novo* genes evolve within a limited evolutionary time scale. Some studies found that protein disorder increases with the age of *de novo* genes in *Lachancea* (Vakirlis et al., 2018), and some other studies suggested *de novo* genes evolved or adapted gradually to avoid disorder (Wilson et al., 2017). While some case studies (Bungard et al., 2017; Lange et al., 2021) and a recent study focusing on *de novo* ORFs (Schmitz et al., 2018) suggested that protein disorder and other properties hardly change after origination. Here, by comparing *de novo* genes originated from different branches, our results supported that protein disorder and other structural properties change little after origin. Interestingly, other properties, such as GC content, evolutionary rate, and expression pattern change with the age of *de novo* genes. Specifically, compared to younger *de novo* genes, older *de novo* genes have higher GC content, slower evolutionary rate, and are more broadly expressed in various tissues. The results also indicated that GC content was not necessarily the cause of high protein disorder in *de novo* genes, as protein disorder hardly changes while GC content decreases with the age of *de novo* genes. The changes in GC content might be contributed by mechanisms such as the gradual optimization of codons, which could potentially increase the translational efficiency of *de novo* genes. We noticed that *de novo* genes have increased strength of purifying selection (decreased ω_na_) with *de novo* gene ages, while the adaptation rates change little, which suggests that older *de novo* genes might be related to more important functions. In agreement with this result, it was reported that, at a larger evolutionary timescale, the number of PPIs and genetic interactions increased gradually with the age of genes (Abrusán, 2013). Taken together, we proposed that, upon origination, *de novo* genes were involved in some molecular functions with certain fitness effects, partly due to the tendency to encode signal proteins or transmembrane proteins (Lange et al., 2021; Rivard et al., 2021; Vakirlis, Acar, et al., 2020). In cases where de novo genes lost their functions, they might be depleted shortly after origination (Palmieri et al., 2014; Zheng & Zhao, 2022). They tend to be in the periphery of cellular networks (Abrusán, 2013), and thus more likely to be tolerated by the host. Due to the weaker selective constraints, *de novo* genes tend to undergo faster sequence evolution, resulting in abundant sequence changes. These changes could potentially happen in regulatory or coding regions and further affect the expression patterns or expression levels of *de novo* genes. The gradual shift of sequence and expression patterns of *de novo* genes might increase their chances of being involved in protein-protein interactions (PPI), genetic interactions or other intermolecular interactions through adaptive evolution (Peng et al., 2022), which can further facilitate the integration of these genes into cellular networks. On the other hand, *de novo* genes might retain certain basic molecular functions by constraining similar overall structures (see section *Most potentially well-folded de novo genes are likely to be born well-folded* in Results) or key sequence motifs along their evolutionary trajectories.

Our study presents a comprehensive analysis of *de novo* genes of various ages, offering a systematic overview on their origination and evolution. While previous investigations in different lineages have employed diverse methodologies (Begun et al., 2007; Carvunis et al., 2012; Heames et al., 2020; Knowles & McLysaght, 2009; Majic & Payne, 2020; Moyers & Zhang, 2016; Ruiz-Orera et al., 2015; Vakirlis et al., 2018; Witt et al., 2019; Wu et al., 2011; Zhao et al., 2014; Zheng & Zhao, 2022), several characteristics of *de novo* genes appear to be consistent across multiple lineages. Notably, *de novo* genes tend to be relatively short in length and exhibit a strong enrichment in testis, displaying biased functions associated with this reproductive organ. Additionally, in mammals, the expression of *de novo* genes in the brain is relatively common (An et al., 2023; Ruiz-Orera et al., 2015; Wu et al., 2011). This is also observed in older *de novo* genes in *Drosophila*, although to a much less degree. Intriguingly, our research highlights the potential for the immune system (Go enrichment P-value=7e-4) to serve as another hotspot for *de novo* gene origination. However, further investigations are required to determine if this pattern holds true in other taxa or lineages. Furthermore, our investigation reveals that *de novo* genes predominantly exhibit a disordered nature, and this characteristic remains stable over the time frame examined. These findings align with the study by Schmitz et al. (Schmitz et al., 2018). In contrast, there are some inconsistencies in previous studies. For example, Carvunis et al. observed in yeast that protein structural disorder increased with gene age (Carvunis et al., 2012), and Wilson et al. observed in yeast and mouse that protein structural disorder decreased with gene age (Wilson et al., 2017). Investigating whether this discrepancy arises from methodological differences or possesses biological relevance warrants further exploration. Our work is the first study to reveal structural conservation for well-folded protein-coding genes using high-accuracy 3D structure modeling, whether this is a conserved pattern awaits future studies in other taxa or lineages. Another major point of contention revolves around the origin of *de novo* genes: whether they arise through neutral processes or are driven by strong selection. Our study on *D. melanogaster,* a species that has a large effective population size, demonstrates that both adaptive and nonadaptive changes play pivotal roles in the slightly accelerated evolution of *de novo* genes after their birth. Previous studies that have distinguished adaptive and nonadaptive rates in *de novo* gene evolution are scarce. Exploring the applicability of our findings to other taxa or lineages, particularly those with smaller effective population sizes like humans, would be a fruitful avenue for future research.

Overall, our study provided a general and easy-to-use pipeline to identify lineage-specific *de novo* gene candidates. We also provided a systematic overview of the protein foldability of *Drosophilinae* lineage-specific *de novo* gene candidates. We proposed that *de novo* genes undergo gradual sequence and functional adaptation without major protein structure changes in *Drosophila* lineage. With recent advances in *de novo* gene detection frameworks, such as those used in this work and in the work of others, e.g., Vikrilis et al., 2020 (Vakirlis, Carvunis, et al., 2020), it would be exciting to identify not only young but slightly older *de novo* genes in other lineages. This would provide an evolutionary framework for comparing *de novo* genes in multiple taxa. With more high-quality genome sequencing data, and transcription and translational data for more related species in the future, we will be able to identify *de novo* genes with higher confidence and uncover more about their emergence and how they were incorporated into the genomes and interactomes.

## Material and Methods

### Identification of de novo gene candidates in D. melanogaster

To infer synteny, we used Progressive Cactus Genome Aligner (Armstrong et al., 2020) to align the genomes of 20 *Drosophilinae* species and 2 *Acalyptratae* species (Figure 1A). To infer homology between different protein-coding genes, we also run all-vs-all *blastp* using all the protein sequences of 20 *Acalyptratae* species along with another 8 *Arthropods* species (Figure 1A). After obtaining the synteny and homology information, we used the following workflow (Figure 1B) to identify possible candidates of *de novo* genes.

1. As mentioned above, we used progressive cactus aligner to align the 20 genomes of species in *Acalyptratae* lineages. The species were assigned different branch numbers according to their separation to *D. melanogaster*, ranging from *Br_1_* to *Br_9_* (Figure 1A). In this step, there were 13798 *D. melanogaster* protein-coding genes aligned in cactus whole genome alignments.
2. For each of the annotated query protein-coding genes in one of the 20 *Acalyptratae* species, we used pairwise *halLiftover* to determine the syntenic region in another *Acalyptratae* species.
3. If the syntenic region is an annotated protein-coding gene, we assign it to the ortholog of the query when the annotated gene has an E-value smaller than 0.05 to the query in the all-vs-all *blastp* analysis. Otherwise if the syntenic region is unannotated, we use *Genewise* (Birney et al., 2004) and *Spaln* (Iwata & Gotoh, 2012) to predict protein-coding potential from the unannotated syntenic region using the query gene as reference. We assign the syntenic hit as an unannotated ortholog if the coding potential was significantly beyond random simulations (see next section, *Random simulations of Geneise/Spaln*, for details), otherwise we assign it as a non-genic hit.
4. For each *D. melanogaster* protein-coding gene, we inferred their annotated/unannotated orthologs and non-genic hits based on the pairwise halLiftover in Step 2 and ortholog identification in Step 3. We were able to identify 1285 potential *de novo* gene candidates due to the presence of non-genic hits in their furthest aligned branches in cactus alignments.
5. We assigned the query as a possible *de novo* gene candidate and inferred the origination branch to be *Br_i_* (i ∈ {0…9}) only when it has: (1) annotated or unannotated orthologs and homologous in branches *Br_i_* (Fig. 1A), (2) non-genic hits in the outgroups of *Br_i_*, and (3) no homologs in the 8 distant *Arthropods* species as shown in Fig. 1A. To this preliminary step, we identified 686 potential *de novo* gene candidates, each with the inferred origination branch as the branch that has the most distant orthologs (*Br_i_* mentioned above and main text). These candidates are then subjected to further searches in all other species in UniprotKB and NCBI representative genomes (see below).
6. We then used *jackhmmer* (S. R. Eddy, 2011) and *blastp* to search against UniProtKB sequence database to further filter out candidates that have homologs in species more distant than the inferred origination branch *Br_i_* as defined in step 5 and main text. To control for the possibility of false positives, which were quite frequent in iterative profile searches, we conducted three independent searches as follows:

i. *blastp* search with E-value cutoff of 0.05.
ii. Iterative *jackhmmer* with options “--incE 1e-5 -E 10”.
iii. Iterative *jackhmmer* with options “--incE 1e-5 -E 10”. To control for false positives, after each iteration, we manually built a hmm profile for the next iteration by removing possible false positives with the best 1 domain E-value larger than 1e-5 (S. Eddy, 1992). For each *de novo* gene candidate, we stopped the search once the search converged or reached 5 iterations.

To further control for possible false positives, for each *de novo* gene candidate, we required a reliable homolog to appear in at least two of the above searches with a E-value cutoff of 0.001. At this, we removed candidates that have reliable homologs in species that are more distant than their inferred origination branch obtained in step 5. The removed candidates, along with representative reliable homologs can be found in *supplementary File S4: reliable blastp jackhmmer and tblastn homologs*.
7. As a final step, we used *tblastn* to search for possible unannotated homologs in species that are more distant to the inferred origination branches as defined in step 5 and main text. First, we used *tblastn* to search against NCBI representative genomes at E-value cutoff of 1.0. We then extracted the DNA sequences of the significant hits with the following command, blastdbcmd -db ref_euk_rep_genomes -entry *RefSeqID* -range *START-END* -strand *STRAND* -out out.fasta -outfmt %f where *`RefSeqID`*, *`STRAND`*, and *`START-END`* defined the locations of the significant hits. We extended the range (*START-END*) to match the size of the query *D. melanogaster* protein-coding genes. To further determine whether the *tblastn* hits were possible homologs, we used *Genewise*/*Spaln* to predict protein coding potential from the significant *tblastn* hits using the query *D. melanogaster* protein-coding gene as reference. We manually examined the *Gewise*/*Spaln* predictions. A significant *tblastn* hit was considered as an unannotated homolog if it met the following criteria:

i. The predicted gene has a canonical start and a stop codon.
ii. The predicted gene has no frameshifts.
iii. The predicted gene has the same number of exons and introns as the query *D. melanogaster* gene or its orthologs in other *Drosophila* genomes.

At the *tblastn* filtering step, we removed candidates that have reliable unannotated homologs in the outgroup species of their inferred origination branches obtained in step 6. The removed candidates, along with representative reliable homologs can be found in *supplementary File S4: reliable blastp jackhmmer and tblastn homologs*.
8. After the above filtering steps (step 6 and 7), we were able to identify 555 *de novo* gene candidates in *D. melanogaster* that are potentially originated within *Drosophilinae* lineage. The full list of the 13968 annotated protein-coding gene in *D. melanogaster*, 13798 aligned in cactus, 1285 potential candidates with non-genic hits, 686 preliminary candidates, and final 555 de novo gene candidates can be found in *supplementary File S5: list of D. melanogaster protein-coding genes in the de novo gene identification workflow*.

For each *de novo* gene candidate identified here, we used the longest protein isoforms as the protein sequence for further analysis.

### Random simulations of Genewise/Spaln

We used random simulations to calculate the random expectations of Genewise/Spaln gene structure predictions. The random simulations were carried out as follows:

1. We simulated situations with different protein lengths, ranging from 15, 20, 25, 50, 75, 100, 200, 500 to 1000.
2. For each protein length (*N*), we ran 10,000 random simulations. For each simulation, we generated a random protein sequence according to the ratio of each amino acid in *D. melanogaster*, and a random DNA sequence with 3*(N+300) base pairs according to the genome-wide GC content of *D. melanogaster* protein-coding DNA sequences. We performed Genewise or Spaln gene structure predictions using the protein as reference and the DNA sequence as target. To assess the results of the predictions, we used the score reported by spliced alignment, as the score includes the penalty for introducing splicing sites. We calculated the mean (Mean) and standard deviation (SD) of the 10,000 random simulations.
3. We fitted Mean and SD as functions of protein length (*N*) respectively, using a two-phase decay function using Lmfit (Newville & Stensitzki, 2018)

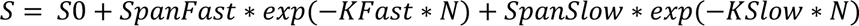

where

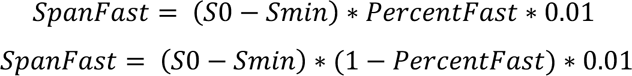

S0 and Smin were the expected maximum and minimum spliced alignment scores. We then can predict the random expectation scores for Genewise/Spaln gene structure prediction score for any protein references with length of N by the two-phase decay function (Figure S8).
4. We computed the probability of the protein reference and DNA target being completely unrelated or non-homologous using one-tailed gaussian distribution with the inferred mean and standard deviations of Genewise/Spaln prediction scores. We inferred the target DNA sequences being potentially genic and homologous to the reference protein at a P-value of 10^−6^.

### Bioinformatic analysis and evolutionary analysis

We used RepeatMasker (Smit AF et al.,) and Dfam (Storer et al., 2021) to search for DNA repeats, including transposable elements (TEs) and simple repeats. We used ORFfinder to detect all putative ORFs that are intergenic and longer than 75 nt (coded for 25 amino acids). We included nested ORFs to better understand possible residue composition of these intergenic putative ORFs. For protein property predictions, we used deepcnf (S. Wang et al., 2016) to predict per residue probability of helix, sheet, coil, structural disorder, solvent accessibility and transmembrane probability. These properties were further normalized by protein length. We used signalp-5.0 (Almagro Armenteros et al., 2019) to determine the probability of a protein containing a signal peptide. Sequence evolutionary rate (ω), adaptation rate (ω_a_), and nonadaptation rates (ω_na_) of *de novo* gene candidates were obtained from a previous study (Peng et al., 2022).

### Gene expression patterns

We downloaded gene expression profiles from FlyAtlas2 (Leader et al., 2018). We converted FPKM to TPM by normalizing FPKM against the summation of all FPKMs as follows:

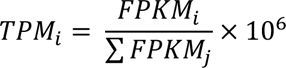

After TPM conversion, we only retained genes with expression levels larger than 0.1 TPM for further analysis. We treated male and female whole-body TPM as male and female expression levels. To describe male specificities of *D. melanogaster* genes, we first calculated Z-score by:

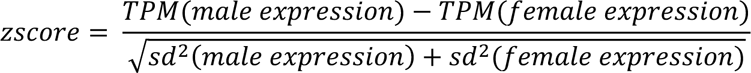

We then calculated a normalized Z-score by:

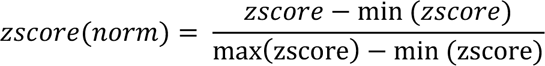

To characterize tissue specificities, we calculated tau values (Yanai et al., 2005) based on the expression profiles of 27 different tissues.

### Structural prediction for de novo gene candidates

We combined AlphaFold2 (Jumper et al., 2021), RoseTTAFold (Baek et al., 2021), trRosetta (J. Yang et al., 2020) and ESMFold (Lin et al., 2023) to predict the foldability of the *de novo* gene candidates. Specifically, for AlphaFold2 predictions, if the *de novo* gene was present in AlphaFold Protein Structure Database (AlphaFold DB), we downloaded the predicted structure from the database, otherwise we ran AlphaFold v2.0.0 scripts (https://github.com/deepmind/alphafold/tree/v2.0.0) to predict the protein structure. For RoseTTAFold predictions, we used the script *run_e2e_ver.sh* to predict the protein structures. For ESMFold, we used the api on the ESM Metagenomic Atlas website (https://esmatlas.com/about#api) to fold the protein sequences. We used the per-residue confidence score (pLDDT) from AlphaFold2, RoseTTAFold, and ESMFold predictions to predict the foldability of the *de novo* genes. For trRosetta predictions, we followed similar procedures as DeepMSA (C. Zhang et al., 2020) and iteratively searched through Uniclust30 (Mirdita et al., 2017), UniRef90 (Suzek et al., 2015), metaclust (Steinegger & Söding, 2018) and tara(Y. Wang et al., 2019) protein sequence databases at default E-value cutoff of 1e-3 to collect homologous or distantly related homologous sequences. Specifically, at the first stage (Uniclust30), we used hhblits (Remmert et al., 2012) to search against Uniclust30. At each stage from stage 2 to stage 4 (UniRef90, metaclust, and tara), we first used jackhmmer (S. R. Eddy, 2011) to search against the database and then used hhblits against a custom database built from the sequences generated by jackhmmer. After each stage of searching, we used MAFFT to construct a multiple sequence alignment (MSA) to ensure that the MSA is of high quality. With the final MSA as input, we generated 100 parallel structural models by trRosetta (J. Yang et al., 2020). To characterize the foldability of the *de novo* genes and the convergence of the predictions, we selected an ensemble of 20 structural models with the lowest potential energies and calculated the average pairwise TM-score (Y. Zhang & Skolnick, 2004) by TM-align (Y. Zhang & Skolnick, 2005).

### Ancestral sequence reconstruction

For each *de novo* gene, we extracted their orthologs and unannotated putative orthologs (if there are any) from the above analysis (see section *Identification of de novo gene candidates in D. melanogaster* in Material and Methods). We used MAFFT-LINSI (Katoh et al., 2002) to align the orthologous protein sequences. An initial guide tree was built based on the phylogeny relationships between the species that had orthologs. We then used the AAML module in PAML 4.9 (Z. Yang, 2007) to reconstruct the most probable ancestral sequences based on the alignments and the initial guide tree. For each branch, we only considered amino acids that had more than 50% coverage among the species in this branch, as the reconstruction can be unreliable at sites with alignment gaps. We further used AlphaFold2 to predict the structures of the most probable ancestral sequences.

### Molecular dynamics simulations

Starting from the predicted structural models of the 19 potentially well-folded *de novo* gene candidates and their most probable ancestral states, we set up the molecular dynamics simulations using the GROMACS-2019 (Abraham et al., 2015) and the Amber ff99SB-ILDN force field (Lindorff-Larsen et al., 2010). For each system, we placed the protein in a dodecahedral box, with a minimum distance between the solute and the box boundary of 1.2 nm. We filled the simulation box with TIP3P water molecules and additional Na^+^ and Cl^−^ ions to neutralize the system and reach the 0.15 M salt concentration. We further minimized the potential energy of the system by the steepest descent method followed by conjugate gradient method until the maximum force was smaller than 200 kJ mol^−1^ nm^−1^. For each simulation system, we performed 200 ns MD simulations. We clustered the conformations from the last 150 ns simulations into 5 clusters by the density peaks algorithm (Rodriguez & Laio, 2014). To obtain structural fold similarity during MD simulations, we used TMalign (Y. Zhang & Skolnick, 2005) to compute the pairwise TM-score of the centers of the 5 clusters. We also referred to the center of the 5 clusters as representative structural models and the center of the top cluster as the MD refined structural model during MD simulations.

### Clustering of testis-biased genes

For each testis-biased gene, we obtained the scaled expression levels at different germline cell development stages, including 1) Early spermatogonia, 2) Late spermatogonia, 3) Early spermatocytes, 4) Late spermatocytes, 5) Early spermatids, and 6) Late spermatids. We further applied principal components analysis (PCA) on the scaled expression dataset to reduce the dimensionality. We then applied a k-means clustering algorithm to cluster these genes using the first three PCs, which could explain 82.0%, 12.2%, and 3.2% of the variants, respectively. The testis-biased genes were finally clustered into four clusters by kmeans clustering method (Figure S5). The sum of squared error of kmeans clustering was also shown in Figure S5.

## Data Availability

Whole genome alignments used in this study are available at 10.6084/m9.figshare.19989395, along with the raw alignments and spliced alignments of *de novo* gene candidates, which include non-coding hits in outgroup species. Code and scripts for this study are available on GitHub: https://github.com/LiZhaoLab/DrosophilaDenovoGene.

## Acknowledgements

We thank members of the Zhao laboratory for helpful discussions, and Nicolas Svetec, Eli Woloshin, and Vivian Yan for critically reading the manuscript.

