## Supplementary figures and tables for "The origin and structural evolution of *de novo* genes in *Drosophila*"

Supplementary figures and tables for
**The origin and structural evolution of *de novo* genes in *Drosophila***

Junhui Peng, Li Zhao
Laboratory of Evolutionary Genetics and Genomics, The Rockefeller University, New York, NY 10065, USA

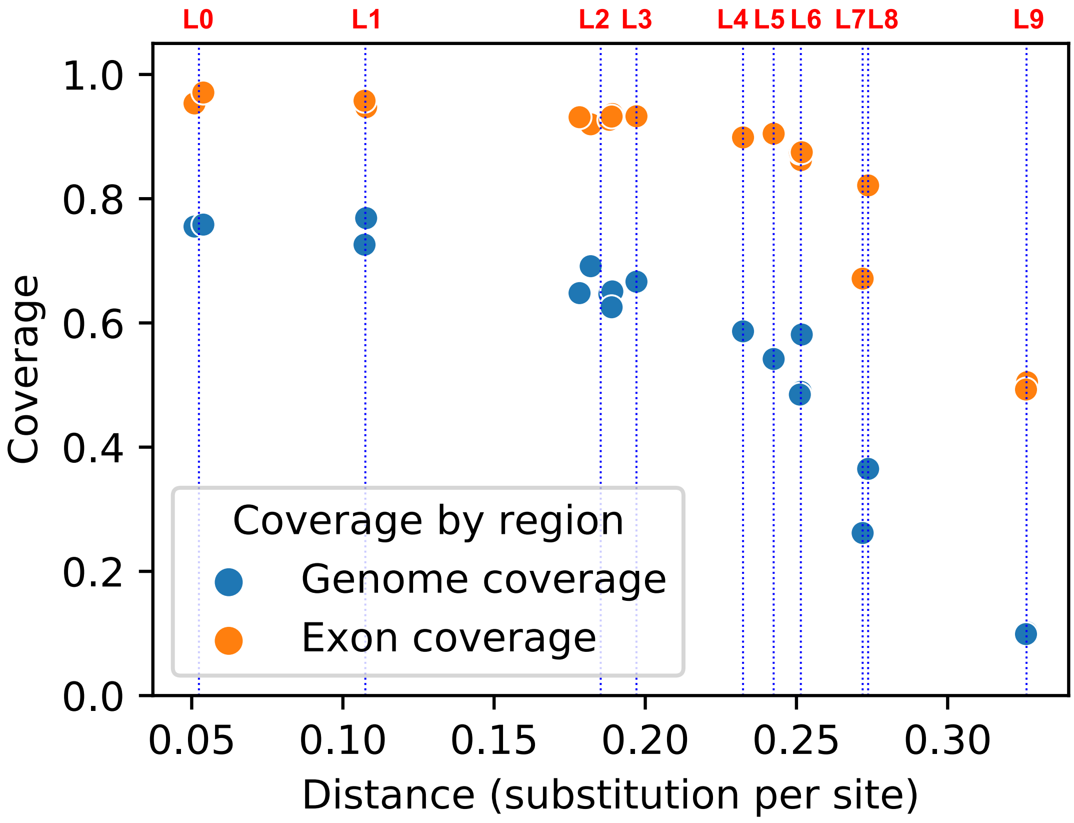


Figure S1. Statistics of progressive cactus alignments.

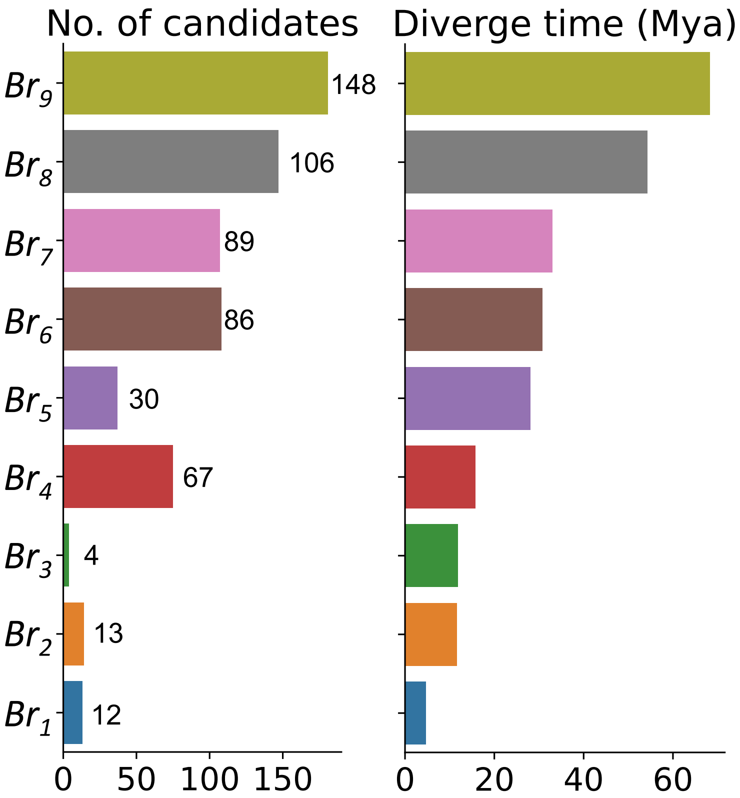


Figure S2. Number of *de novo* gene candidates identified in each branch. The number generally correlated with the divergence time between *D. melanogaster* and each branch.

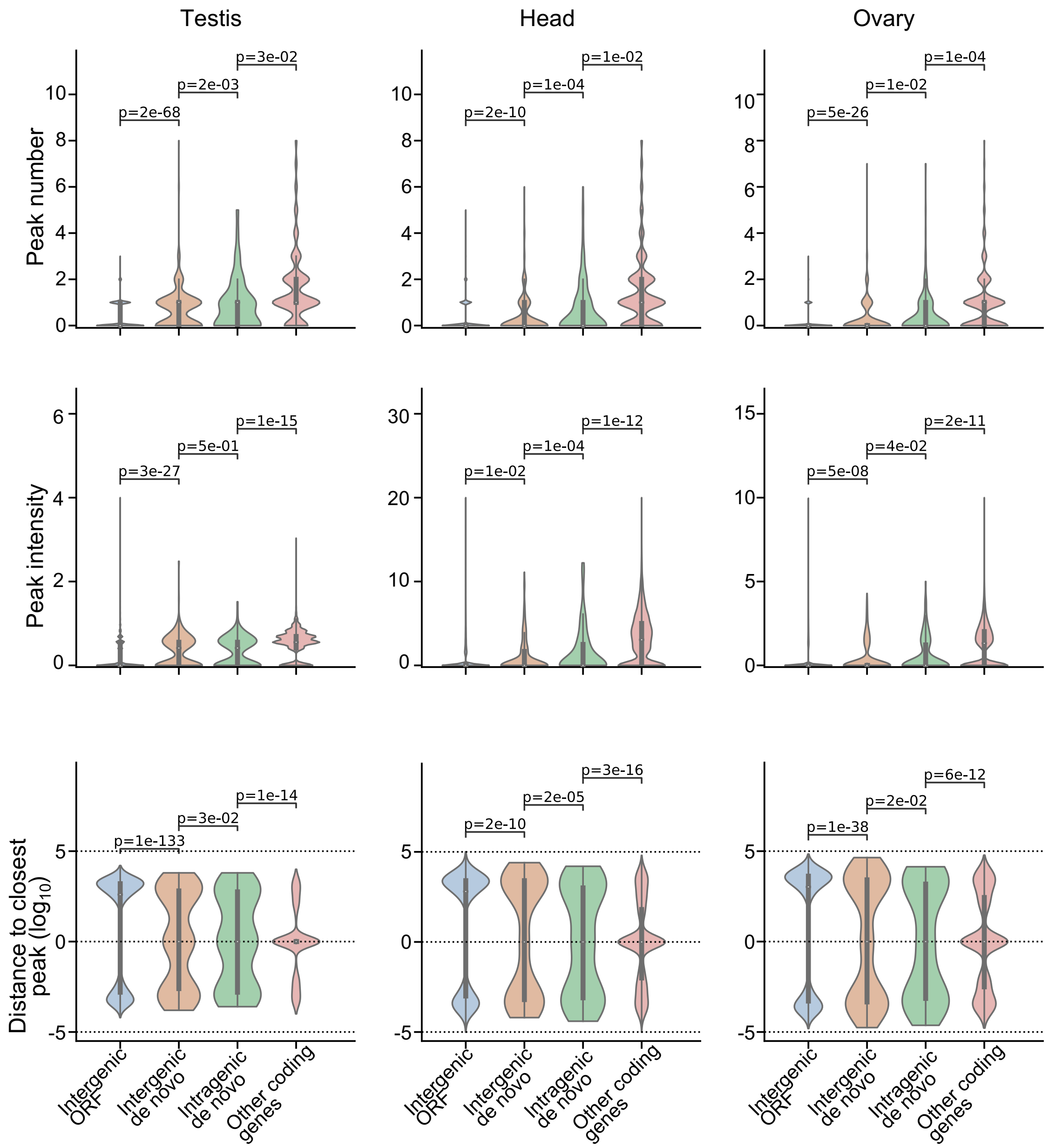


Figure S3. ATAC-seq peaks, intensities, and distances for de novo gene candidates in testis, head, and ovary.

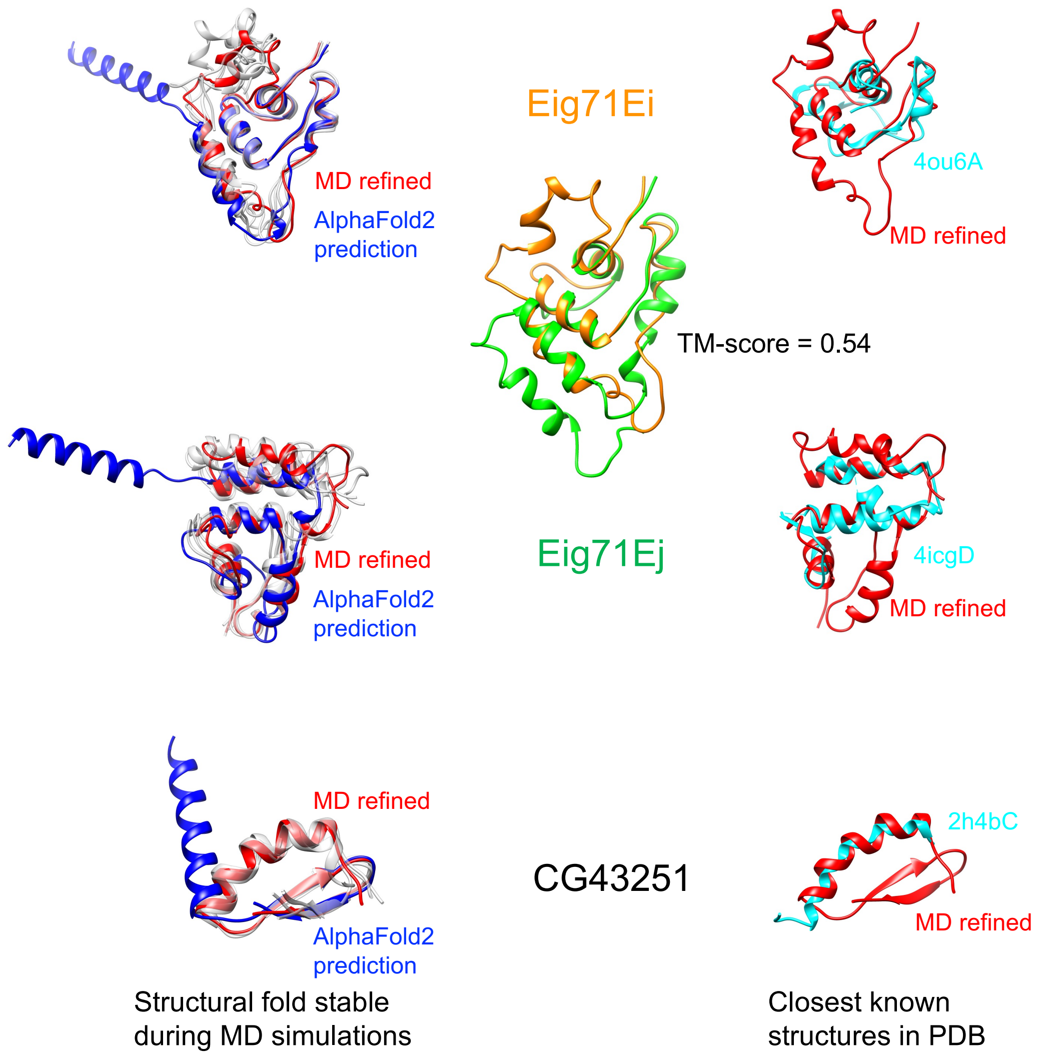


Figure S4. Eig71Ei, Eig71Ej, and CG43251 might adopt new structural folds. Left panel: Eig71Ei, Eig71Ej, and CG43251 remained similar structural folds during MD simulations. Right panel: the structures in PDB with the largest TM-scores to Eig71Ei, Eig71Ej, and CG43251 were superimposed to their MD refined structural models. All the three largest TM-scores were smaller than 0.5 (Table S1). Eig71Ei and Eig71Ej are paralogs and share similar structural folds with TM-score of 0.54 (inserted panel).

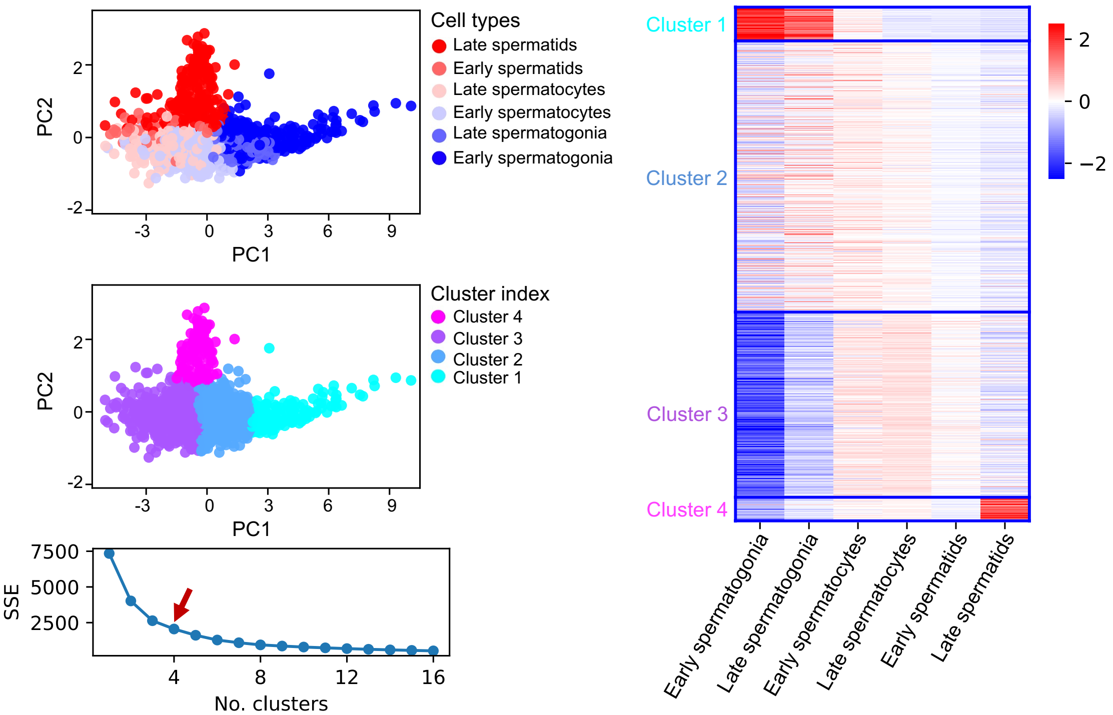


Figure S5. Clustering of all *D. melanogaster* testis-biased genes. The sum of squared error (SSE) as a function of the number of clusters was shown in the bottom left panel.

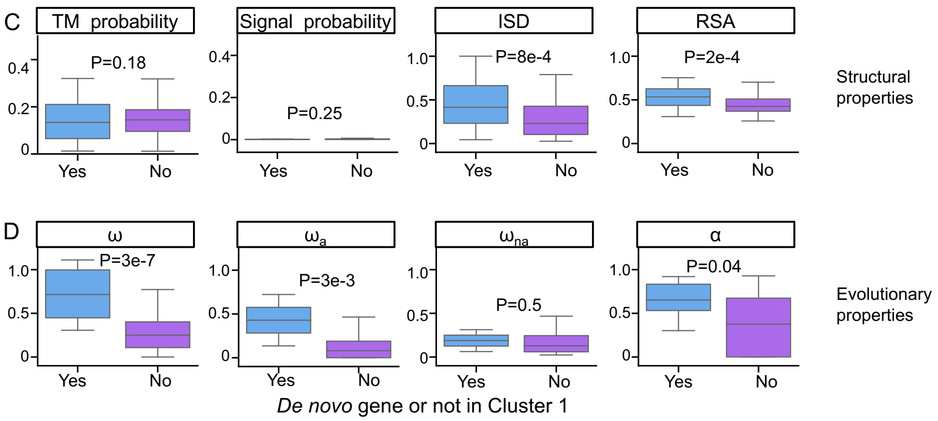


Figure S6. Comparison of testis-biased *de novo* genes (cyan) and non-*de novo* genes (purple) in cluster #1. For clustering analysis, see Figure 5 for detail. In cluster #1, testis biased de novo genes are more disordered (ISD panel) and exposed (RSA panel). These de novo genes also evolve faster ($\omega$ panel) with higher adaptation rates ($\omega_a$ panel).

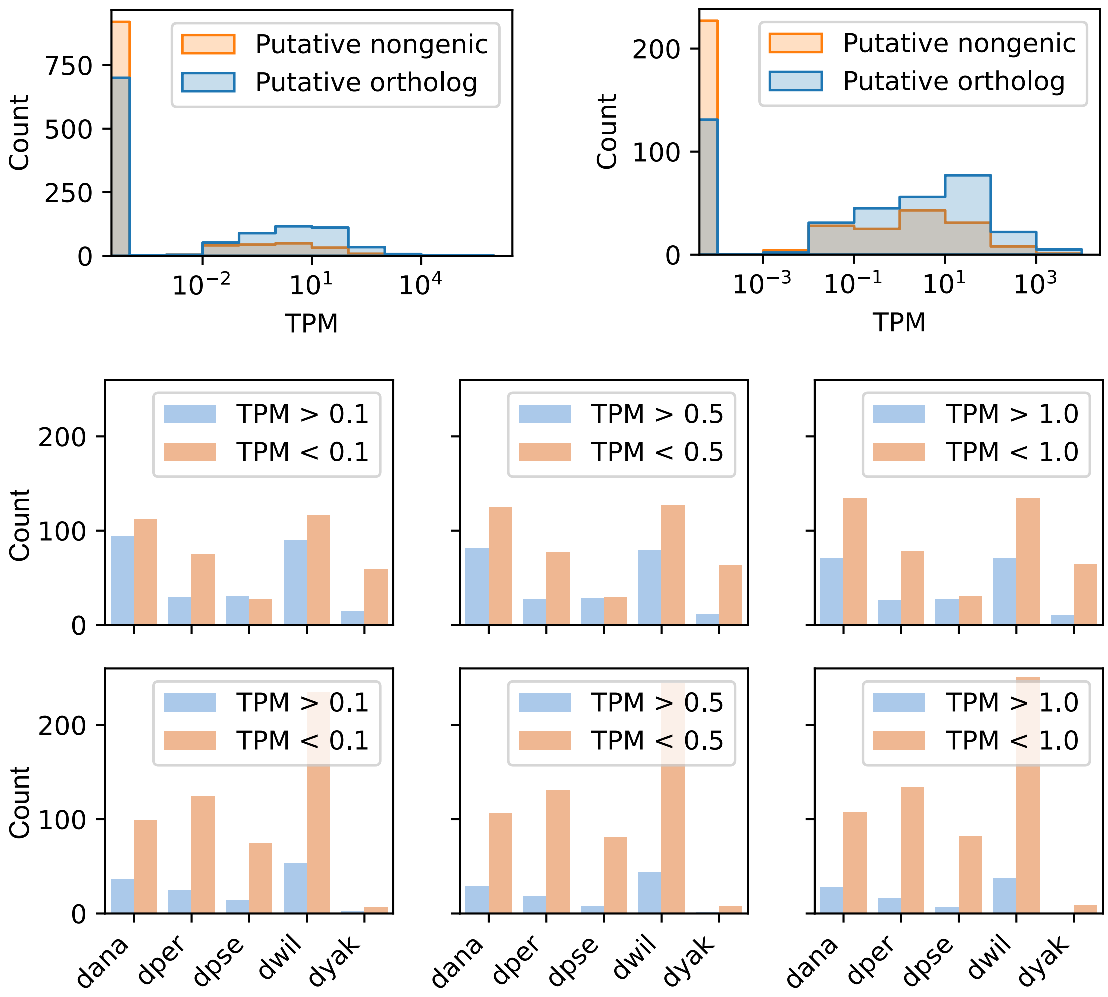


Figure S7. RNA-seq support of the unannotated putative orthologs of some de novo gene candidates in dyak, dana, dper, dpse, dwil.

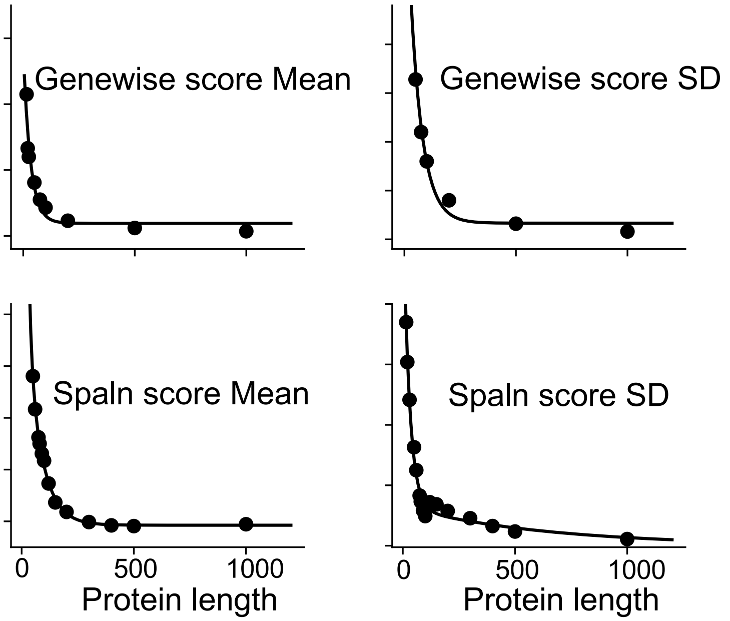


Figure S8. Random simulations of Genewise/Spaln. We used two-phase decay function to fit the mean (left panel) and standard error (right panel) of spliced align score (Genewise, top panel, and Spaln, bottom panel).

Table S1. Proportions of optimal codons in *de novo* genes and other annotated protein-coding genes in *D. melanogaster*. The median values of the proportions were listed in the P(optimal, *de novo*) and P(optimal, other) columns. *De novo* genes show significant less optimal codon usage compared to other annotated protein coding genes. The P-value was computed using *scipy.stats.ttest_ind* module with option *alternative="less"*. was shown in the P(t-test) column. For most of the amino acids, the proportion of optimal codons show significant positive correlation with the origination branches as shown by the P-values of Spearmanr and Kendalltau rank correlation test.

| Amino Acid | P(optimal, *de novo*) | P(optimal, other) | P(t-test) | P(Spearmanr) | P(Kendalltau) |
|---|---|---|---|---|---|
| A | 0.32 | 0.45 | 9.2E-70 | 4.4E-07 | 5.3E-07 |
| C | 0.62 | 0.73 | 3.1E-25 | 5.9E-02 | 5.8E-02 |
| D | 0.38 | 0.46 | 1.3E-16 | 5.8E-01 | 5.6E-01 |
| E | 0.50 | 0.68 | 7.1E-95 | 6E-07 | 9.9E-07 |
| F | 0.50 | 0.63 | 1.6E-18 | 2.5E-03 | 2.6E-03 |
| G | 0.28 | 0.42 | 2.2E-59 | 2.2E-07 | 2.2E-07 |
| H | 0.50 | 0.60 | 5.2E-31 | 3E-04 | 3.8E-04 |
| I | 0.33 | 0.48 | 2.6E-22 | 1.7E-03 | 1.9E-03 |
| K | 0.53 | 0.71 | 1.5E-52 | 1.2E-01 | 1.2E-01 |
| L | 0.25 | 0.42 | 5.6E-94 | 4.5E-04 | 4.5E-04 |
| M | 1.00 | 1.00 | nan | nan | nan |
| N | 0.50 | 0.55 | 5.6E-14 | 3.7E-01 | 3.9E-01 |
| P | 0.23 | 0.33 | 3E-40 | 2.6E-03 | 3.1E-03 |
| Q | 0.50 | 0.71 | 8.2E-71 | 5.9E-03 | 5.9E-03 |
| R | 0.14 | 0.30 | 6.8E-70 | 5E-07 | 9.7E-07 |
| S | 0.19 | 0.24 | 1.5E-18 | 1.8E-02 | 1.7E-02 |
| T | 0.25 | 0.38 | 1.8E-38 | 5E-05 | 5.7E-05 |
| V | 0.33 | 0.47 | 3.9E-70 | 3.2E-04 | 3.7E-04 |
| W | 1.00 | 1.00 | nan | nan | nan |
| Y | 0.50 | 0.64 | 1.3E-09 | 1.5E-02 | 1.4E-02 |

Table S2. Detailed information of potentially well folded *de novo* gene candidates. Candidates with TM-score to structures in PDB smaller than 0.5 were highlighted in red.

| FBID | pLDDT | Length | Origination lineage | pLDDT (Anc) | TM-score (ToPDB) | Similar fold in PDB | Sequence identity | Gene |
|---|---|---|---|---|---|---|---|---|
| **FBgn0004593** | 0.89 | 98 | L7 | 0.92 | 0.40 | 4ou6A | 0.087 | Eig71Ef |
| **FBgn0014850** | 0.92 | 98 | L7 | 0.92 | 0.45 | 4icgD | 0.053 | Eig71Ej |
| **FBgn0262896** | 0.80 | 39 | L5 | 0.70 | 0.49 | 2h4bC | 0.158 | CG43251 |
| **FBgn0260967** | 0.90 | 280 | L8 | 0.91 | 0.56 | 6xgxB | 0.075 | CG42590 |
| **FBgn0265834** | 0.85 | 153 | L4 | 0.84 | 0.68 | 1u89A | 0.066 | CG44623 |
| **FBgn0261580** | 0.88 | 137 | L6 | 0.78 | 0.60 | 4pr9F | 0.066 | CG42690 |
| **FBgn0261587** | 0.86 | 139 | L7 | 0.86 | 0.59 | 5fjeB | 0.022 | CG42697 |
| **FBgn0263250** | 0.87 | 127 | L3 | 0.88 | 0.61 | 1x91A | 0.023 | CG43393 |
| **FBgn0261581** | 0.85 | 140 | L6 | 0.78 | 0.61 | 6q6bD | 0.028 | CG42691 |
| **FBgn0262819** | 0.91 | 114 | L6 | 0.77 | 0.62 | 5figF | 0.062 | CG43190 |
| **FBgn0265046** | 0.91 | 118 | L6 | 0.81 | 0.64 | 5fjdB | 0.076 | CG44163 |
| **FBgn0052192** | 0.87 | 136 | L3 | 0.86 | 0.64 | 5fjdB | 0.074 | CG32192 |
| **FBgn0037042** | 0.93 | 195 | L8 | 0.92 | 0.65 | 7jh6B | 0.093 | CG12984 |
| **FBgn0264748** | 0.93 | 374 | L6 | 0.90 | 0.66 | 1yrgB | 0.096 | CG44006 |
| **FBgn0264747** | 0.92 | 370 | L6 | 0.90 | 0.66 | 1yrgB | 0.138 | CG44005 |
| **FBgn0264746** | 0.92 | 368 | L6 | 0.90 | 0.67 | 6obnC | 0.133 | CG44004 |
| **FBgn0262480** | 0.89 | 126 | L5 | 0.67 | 0.68 | 6q58D | 0.064 | CG43070 |
| **FBgn0262824** | 0.85 | 138 | L6 | 0.78 | 0.70 | 1u89A | 0.044 | CG43195 |
| **FBgn0263647** | 0.92 | 122 | L5 | 0.91 | 0.76 | 5me8A | 0.057 | CG43638 |

Table S3. MD simulations of 19 potentially well-folded *de novo* gene candidates. Details of the calculation of structural similarity during MD simulations can be found in Material and Methods.

| FBID | Name | RMSD_FL | RMSD_CORE | TM-score |
|---|---|---|---|---|
| **FBgn0037042** | CG12984 | 1.24 | 1.24 | 0.95 |
| **FBgn0264748** | CG44006 | 1.92 | 1.41 | 0.96 |
| **FBgn0264747** | CG44005 | 1.46 | 1.37 | 0.97 |
| **FBgn0014850** | Eig71Ej | 1.55 | 1.51 | 0.87 |
| **FBgn0263647** | CG43638 | 1.06 | 1.06 | 0.95 |
| **FBgn0264746** | CG44004 | 1.49 | 1.29 | 0.97 |
| **FBgn0262819** | CG43190 | 1.04 | 1.04 | 0.95 |
| **FBgn0265046** | CG44163 | 1.17 | 1.12 | 0.94 |
| **FBgn0260967** | CG42590 | 0.99 | 0.99 | 0.98 |
| **FBgn0004593** | Eig71Ef | 1.77 | 1.30 | 0.89 |
| **FBgn0262480** | CG43070 | 0.72 | 0.72 | 0.97 |
| **FBgn0261580** | CG42690 | 2.57 | 1.89 | 0.85 |
| **FBgn0052192** | CG32192 | 1.99 | 1.60 | 0.91 |
| **FBgn0263250** | CG43393 | 0.82 | 0.82 | 0.97 |
| **FBgn0261587** | CG42697 | 1.14 | 0.99 | 0.96 |
| **FBgn0265834** | CG44623 | 2.55 | 1.60 | 0.91 |
| **FBgn0261581** | CG42691 | 1.67 | 1.50 | 0.91 |
| **FBgn0262824** | CG43195 | 1.40 | 1.36 | 0.92 |
| **FBgn0262896** | CG43251 | 1.28 | 1.02 | 0.81 |
